## supplement for "Geodesics to Characterize the Phylogenetic Landscape"

#### Contents


S1 Primate Dataset $\boldsymbol{D_1}$ . . . . . S2

S1.1 Three trees chosen to illustrate pathtrees . . . . . S2

S1.2 Best trees found by PATHTREES, PAUP*, and REVBAYES . . . . . S2

S2 Snake Dataset $\boldsymbol{D_2}$ : Best Trees Found by PATHTREES, PAUP*, and RAXML . . . . . S2

S3 Effect of Interpolation Methods on Visualization . . . . . S4

S4 Validating the MDS Visualization . . . . . S6

S4.1 Validating for primate dataset $\boldsymbol{D_1}$ . . . . . S6

S4.2 Validating for snake dataset $\boldsymbol{D_2}$ . . . . . S7

S5 REVBAYES Script to Generate a Chain of Trees . . . . . S8

### S1 Primate Dataset $D_1$

#### S1.1 Three trees chosen to illustrate pathtrees

Fig. S1 shows the three trees that we have selected in Fig. 3 to give an example of pathtrees.

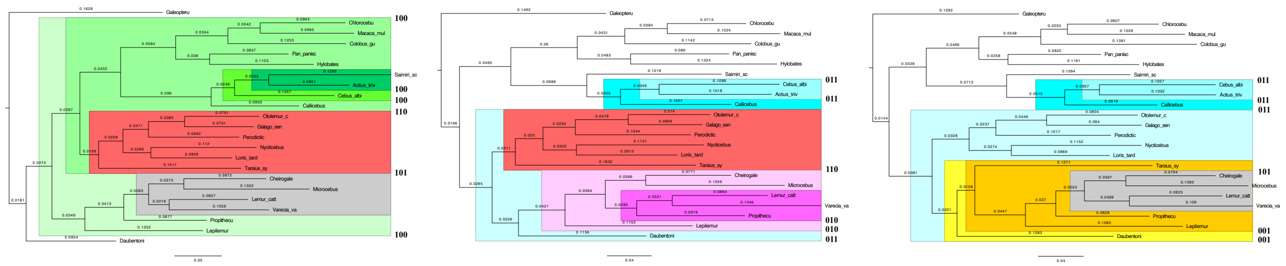

(a) tree ▲ (b) tree ▲ (c) tree ▲

Figure S1. The three trees used in Fig. 3. Colors explain the similarities and differences of the splits in trees. Blue spectrum (011) shows subtrees in trees (b) and (c) and not in tree (a). Red color (110) shows the subtree in trees (a) and (b) and not in the tree (c). Grey color (101) shows the subtree in trees (a) and (c) and not in the tree (b). Green (100), pink (010), and yellow (001) spectrums show the subtrees just in tree (a), tree (b), and tree (c), respectively.

#### S1.2 Best trees found by PATHTREES, PAUP*, and REVBAYES

For the first dataset, we compared our  tree with the maximum posterior tree (MAP) of REVBAYES and the best tree found by PAUP*. As shown in Fig. S2,  our best tree and  PAUP* tree are the same, whereas  MAP differs from both  by two splits. These trees are shown on the likelihood landscape in Fig. 5 and Fig. 6.

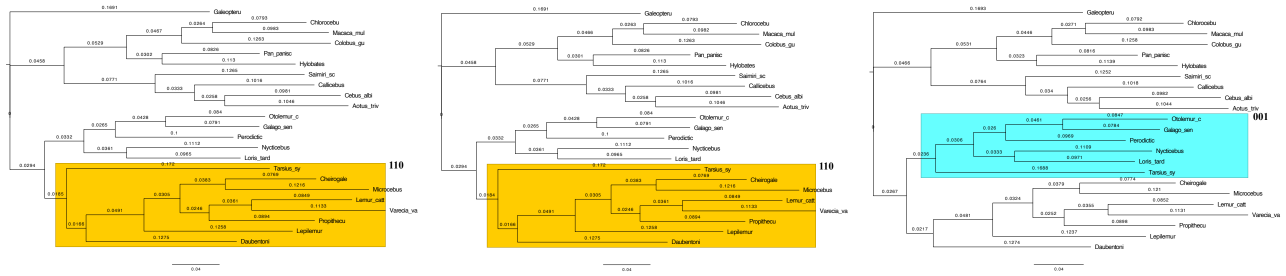

(a) Best tree by PATHTREES (b) Best tree by PAUP* (c) Best tree by REVBAYES

Figure S2. Best trees observed by PATHTREES (a), PAUP* (b), and REVBAYES (c) for the first dataset. Yellow color (110) shows subtrees in the best trees found by PATHTREES and PAUP* and not in the best tree of REVBAYES. Blue color (001) shows the subtrees just in the best tree of REVBAYES.

### S2 Snake Dataset $D_2$ : Best Trees Found by PATHTREES, PAUP*, and RAxML

For the second dataset, we compared our best tree with the best trees found by PAUP* and RAxML. Fig. S3 shows these three trees with colors showing the similarities and differences of the splits in trees.

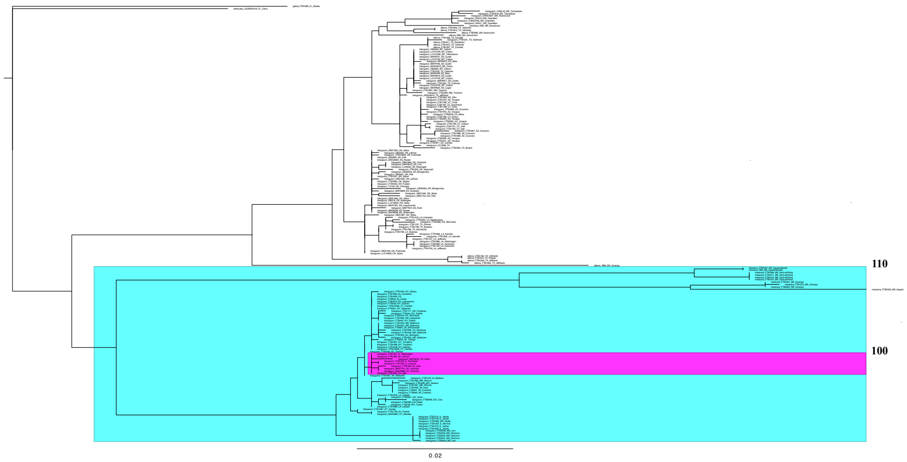


(a) Best tree by Pathtrees

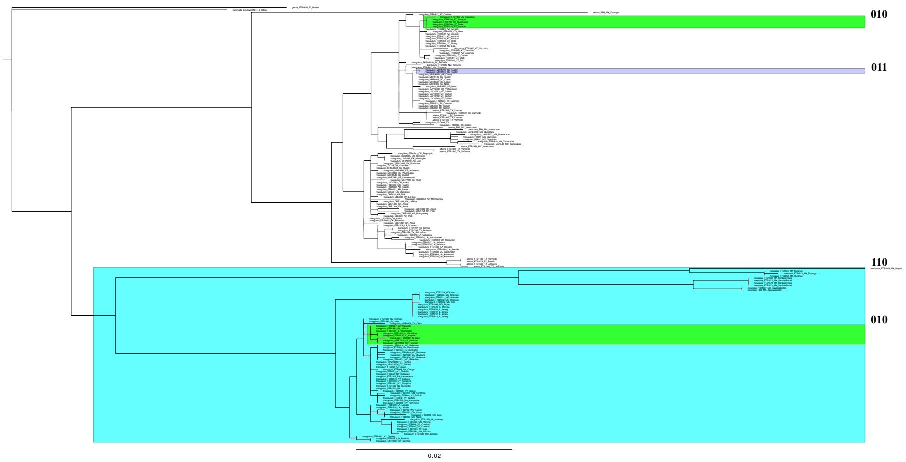


(b) Best tree by Paup*

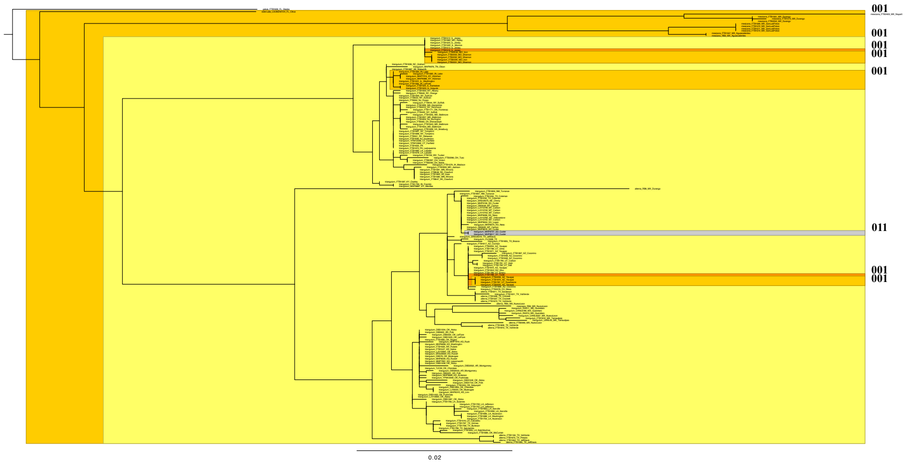


(c) Best tree by RAxML

Figure S3. Best trees observed by Pathtrees (a), Paup* (b), and RAxML (c) for the second dataset. Blue color (110) shows the subtree in the best trees found by Pathtrees and Paup* and not in the best tree of RAxML. Purple color (011) shows the subtree in the best trees found by Paup* and RAxML and not in the best tree of Pathtrees. Pink color (100), green color (010), and yellow spectrum (001) show the subtrees just in the best tree of Pathtrees, Paup*, and RAxML, respectively.

#### S3 Effect of Interpolation Methods on Visualization

Our package PATHTREES uses two different interpolation methods for the likelihood contour and surface. The default interpolation method is the RBF thin-plate spline with smoothness parameter of $s = 1e - 10$. The cubic spline interpolation can be used by setting the option `-interpolate cubic`, and the RBF thin-plate spline with any value of smoothness $s$ can be used by setting the option `-interpolate rbf,s`. The thin-plate spline delivers surfaces that are less noisy. For example, comparing Fig. 7 (thin-plate spline) an Fig. S4 (cubic spline), the overall impression of the contour and surface features are similar but the cubic spline interpolation on the top row shows more extreme peaks than the thin-plate spline interpolation, and the distribution of the range of observed log-likelihood values on the color bar shows this clearly. Therefore, the RBF thin-plate spline interpolation gives a better impression of the true likelihood surface.

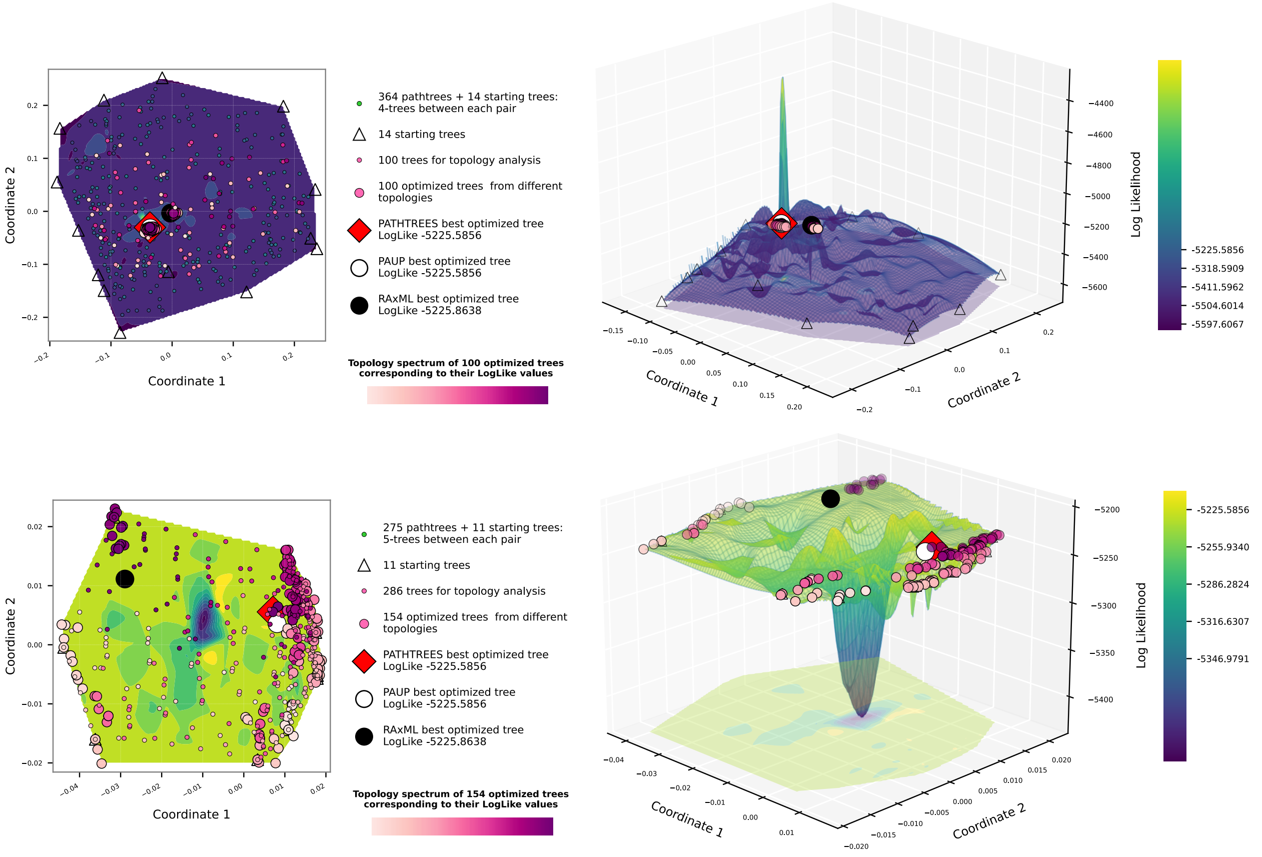


Figure S4. Contour and surface plots of PATHTREES for the dataset $D_2$ , using MDS and cubic spline interpolation defined by the weighted Robinson-Foulds distance metric. First row: the first iteration of PATHTREES using 14 starting trees. Second row: the second iteration of PATHTREES displaying the treespace after zooming inside the convex hull of 100 optimized trees from the first iteration.

Interpolation can fail if many trees are mapped close together in the MDS plot but have sufficiently different likelihoods. RBF will fail in such situations when the surface is forced through the observed data points or will give a singular matrix error (Fig. S5 bottom row). As a solution, increasing the smoothness parameter of the RBF interpolation from zero to a small value, such as $10^{-10}$ can lead to an acceptable visualization (Fig. S5 middle row) compared to the cubic spline interpolation (Fig. S5 top row). The cubic spline interpolation exaggerates the surface compared to the RBF thin-plate spline interpolation.

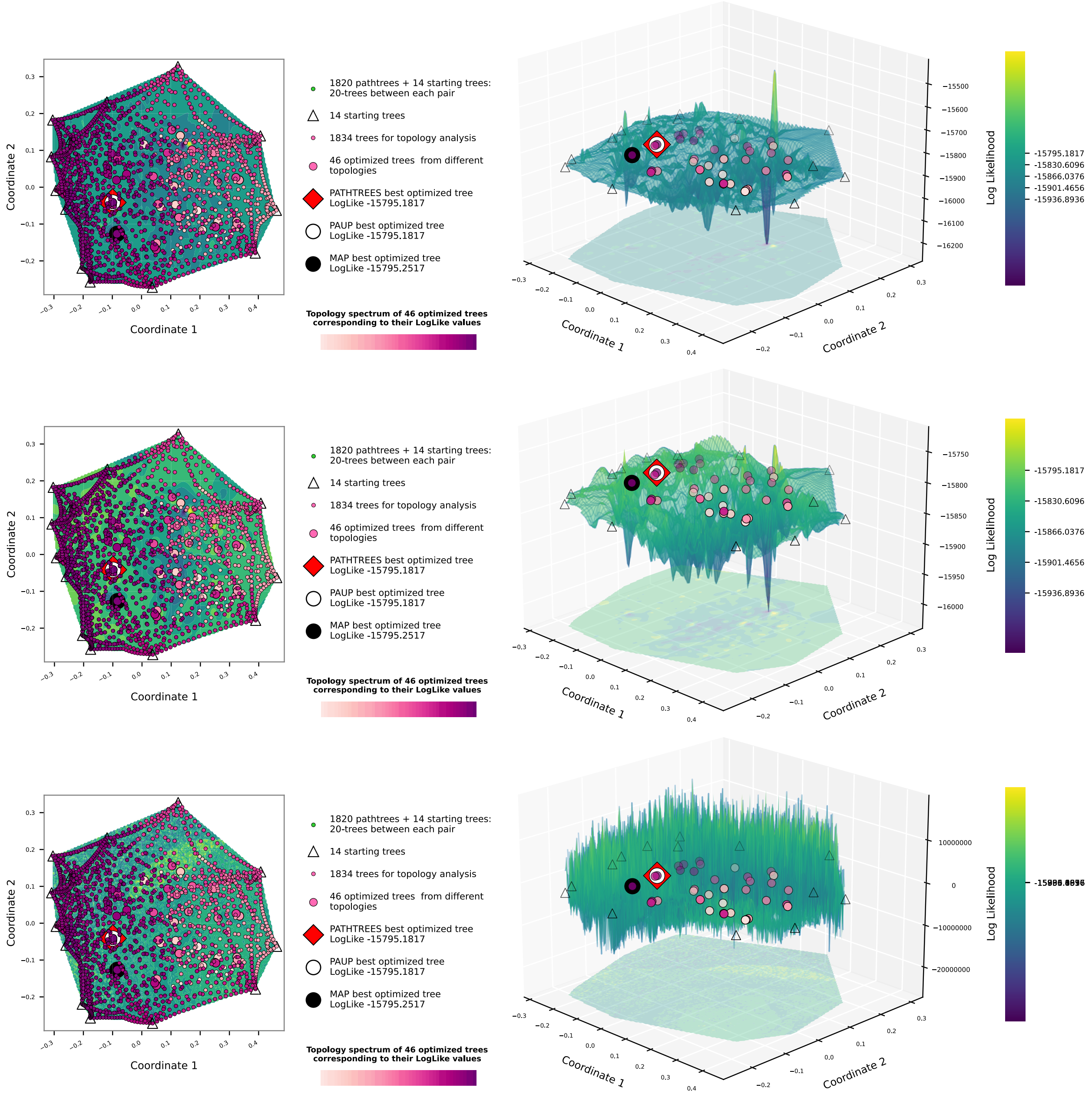


Figure S5. Contour and surface plots of Pathtrees for dataset $D_1$ by generating 20 pathtrees per anchor tree pair, using MDS, and (top row) cubic spline interpolation, (middle row) thin-plate spline with smoothness parameter of $s = 1e - 10$, (bottom row) thin-plate spline with smoothness parameter of $s = 0.0$, defined by the weighted Robinson-Foulds distance metric.

We use MDS to place all trees onto a 2-D plane and then use the likelihood of the trees to interpolate the likelihood tree landscape. This visualization of the MDS plane may differ dependent on the trees used and once in a while may deliver plots that are difficult to interpret; Fig. S6 shows such a plot where the viewpoint of the plane shifted to be perpendicular to our original convex hull: we look at the convex hull sideways. In such cases, changing the number of optimized trees or the number of pathtrees between the anchor trees may help to remedy the view; for example, Fig. 7 (top row) shows an adequate visualization by using fewer number of optimized trees compared to Fig. S6.

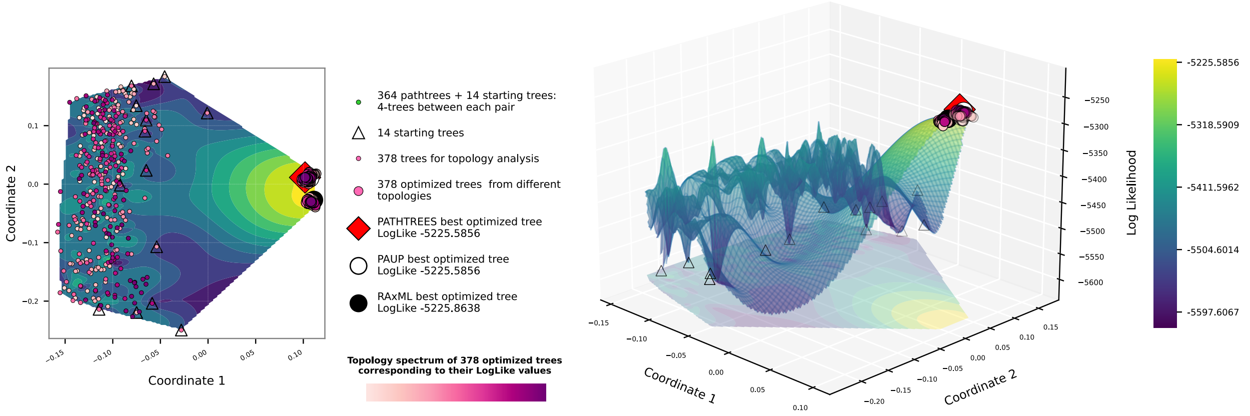


Figure S6. Contour and surface plots of PATHTREES for dataset $D_2$ , using MDS and thin-plate spline interpolation defined by the weighted Robinson-Foulds distance metric. Four trees were generated on the geodesic of each pair of 14 starting trees (364 pathtrees). All 364 + 14 trees were selected to be optimized. All trees have different topologies.

### S4 Validating the MDS Visualization

#### S4.1 Validating for primate dataset $D_1$

Fig. S7 shows the Shepard plot of MDS distances vs. the original dissimilarities for Fig. 5 on bottom row.

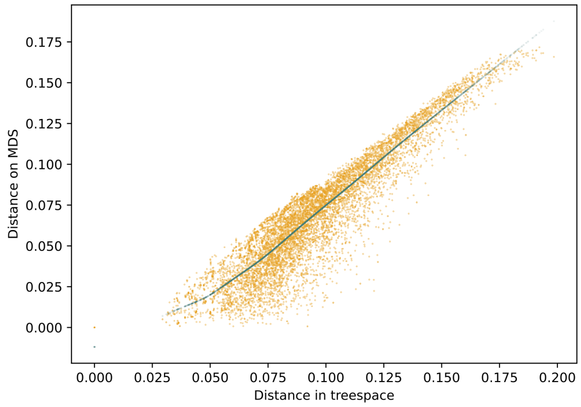


Figure S7. Shepard diagrams of Fig. 5, bottom row, showing the BHV distances versus the MDS distances

We computed the correlation measures Pearson r, Spearman rho, and Kendall tau between the original distances and the MDS distances:

Pearson's r = 0.9236859153718908

Spearman's rho = 0.9071808059933395

Kendall's tau = 0.7517609707944715

It can be seen that the correlation values of Pearson and Spearman are roughly the same, where the value of Kendall correlation coefficient is less than that of others. All these correlation methods are correct in terms of their values. Usually, Spearman's correlation value is closer to the Pearson's value than Kendall's value, and Kendall's value is less that the other two.

##### S4.2 Validating for snake dataset $D_2$

Here, for both iterations of Fig. 7, the Shepard plots of MDS distances versus the real distances are shown in Fig. S8 and the correlation measures Pearson r, Spearman rho, and Kendall tau are computed:

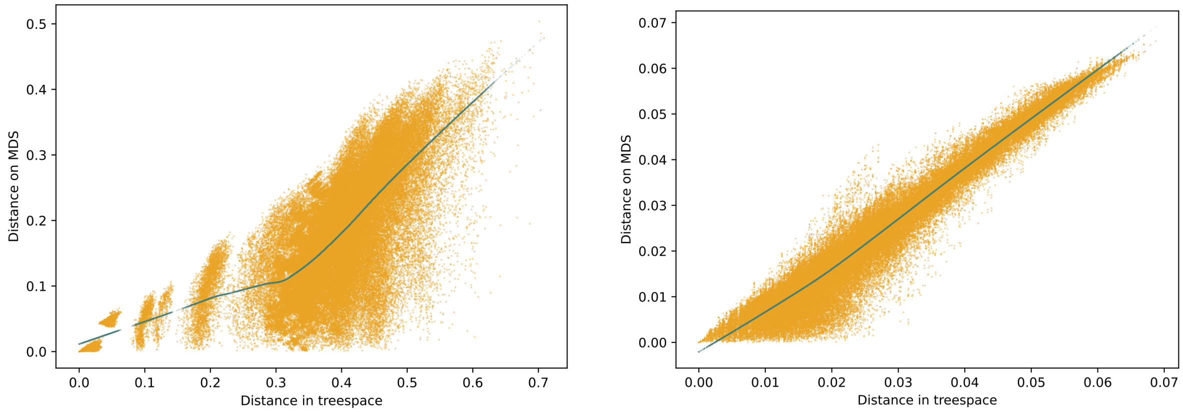


(a) Shepard diagram of the first iteration

(b) Shepard diagram of the second iteration

Figure S8. Shepard diagrams of two iterations in Fig. 7 , showing the BHV distances versus the MDS distances

First iteration:

Pearson's r = 0.6855968829946086

Spearman's rho = 0.6954018788444843

Kendall's tau = 0.518946682776664

Second iteration:

Pearson's r = 0.9702223343002814

Spearman's rho = 0.9573847371609923

Kendall's tau = 0.832968379176889

Since the figure iteration covers a much larger area of trees than the second iteration, the correlations measures are less than those in the second iteration.

#### S5 RevBayes Script to Generate a Chain of Trees

This script is a shortened version of the tutorial page (Höhna et al. 2017)

```
################################################################################
#
# RevBayes Example: Bayesian inference of phylogeny using a Jukes-Cantor
#             substitution model on a single gene.
#
# authors: Sebastian Hoehna, Michael Landis, and Tracy A. Heath
# modified by Peter Beerli 2021
#
################################################################################


### Read in sequence data for both genes
data = readDiscreteCharacterData("data/primates_and_galeopterus_cytb.nex")

# Get some useful variables from the data. We need these later on.
n_species <- data.ntaxa()
n_branches <- 2 * n_species - 3
taxa <- data.taxa()


mvi = 1
mni = 1


#######################
# Substitution Model #
#######################

# create a constant variable for the rate matrix
Q <- fnJC(4)


###############
```

```
# Tree model #
##############

out_group = clade("Galeopterus_variegatus")
# Prior distribution on the tree topology
topology ~ dnUniformTopology(taxa, outgroup=out_group)
moves[mvi++] = mvNNI(topology, weight=5.0)
moves[mvi++] = mvSPR(topology, weight=1.0)

# Branch length prior
for (i in 1:n_branches) {
    bl[i] ~ dnExponential(10.0)
    moves[mvi++] = mvScale(bl[i])
}

TL := sum(bl)

psi := treeAssembly(topology, bl)



###################
# PhyloCTMC Model #
###################

# the sequence evolution model
seq ~ dnPhyloCTMC(tree=psi, Q=Q, type="DNA")

# attach the data
seq.clamp(data)


############
# Analysis #
############
```

```
mymodel = model(psi)

# add monitors
monitors[mni++] = mnScreen(TL, printgen=1000)
monitors[mni++] = mnFile(psi, filename="output/primates_cytb_JC.trees", printgen=10)
monitors[mni++] = mnModel(filename="output/primates_cytb_JC.log", printgen=10)

# run the analysis
mymcmc = mcmc(mymodel, moves, monitors)
#mymcmc.burnin(10000,200)
mymcmc.run(generations=500000)



####################
# Post processing #
####################

# Now, we will analyze the tree output.
# Let us start by reading in the tree trace
treetrace = readTreeTrace("output/primates_cytb_JC.trees", treetype="non-clock")
# and then get the MAP tree
map_tree = mapTree(treetrace,"output/primates_cytb_JC_MAP.tree")


# you may want to quit RevBayes now
q()
```
